# How the healthy ageing brain supports semantic binding during language comprehension

**DOI:** 10.1101/2021.01.15.426707

**Authors:** Roksana Markiewicz, Katrien Segaert, Ali Mazaheri

## Abstract

Semantic binding refers to constructing complex meaning based on elementary building blocks. Using EEG, we investigated the age-related changes in modulations of oscillatory brain activity supporting lexical retrieval and semantic binding. Young and older adult participants were visually presented two-word phrases, which for the first word revealed a lexical retrieval signature (e.g. *swift* vs. *swrfeq*) and for the second word revealed a semantic binding signature (e.g. *horse* in a semantic binding “swift *horse*” vs. no binding “swrfeq *horse*” context). The oscillatory brain activity associated with lexical retrieval as well as semantic binding significantly differed between healthy older and young adults. Specifically for lexical retrieval, we found that different age groups exhibited opposite patterns of theta and alpha modulation, which as a combined picture suggest that lexical retrieval is associated with different and delayed signatures in older compared to young adults. For semantic binding, in young adults we found a signature in the low-beta range centred around the target word onset (i.e. a smaller low-beta *in*crease for binding relative to no binding), while in healthy older adults we found an opposite binding signature about ~500ms later in the low- and high-beta range (i.e. a smaller low- and high-beta *de*crease for binding relative to no binding). The novel finding of a different and delayed oscillatory signature for semantic binding in healthy older adults reflects that the integration of word meaning into the semantic context takes longer and relies on different mechanisms in healthy older compared to young adults.

## Introduction

Healthy ageing is accompanied by decline across a number of cognitive domains, such as your memory for events and the speed with which you process information (Salthouse, 1996; Waters & Caplan, 2005). Language is a crucial aspect of cognition, but the picture of how ageing affects language is a complex one. Older adults get better at some aspects of language, such as knowing more words (Brysbaert et al., 2016), while other skills clearly deteriorate, for example, accessing all the word-related information you need for production (Hardy et al., 2020; Segaert, et al., 2018). At the same time, many other language abilities, including sentence comprehension, appear relatively unchanged by healthy ageing (Peelle, 2019; Shafto & Tyler, 2014). For example, sentence comprehension performance has been demonstrated to be comparable between older and young participants, unless the stimuli are presented at a rapid rate (Tun, 1998; Wingfield et al., 2003) or with background noise (Tun, 1998). The complex behavioural picture for language function is difficult to reconcile with the widespread structural decline in language-relevant brain regions (Antonenko et al., 2013). Even when language performance appears unchanged in older adults, it is likely supported by different functional neural processes from those in young adults (Peelle, 2019). The aim of the current electroencephalography (EEG) study is to investigate the differences between healthy older and young adults in the neural processes involved in semantic comprehension.

When we combine words, the meaning of an individual word (e.g. flat) can be altered by the meaning of a following word (e.g. flat tire vs. flat note) such that the combined meaning is more than the mere sum of its parts (Hagoort et al., 2009; Keenan, 1979). This illustrates the unique and expressive power of language: we have the ability to combine words in novel ways to create sentences. In other words, language users construct complex meaning from more elementary semantic building blocks (Hagoort et al., 2009; Hagoort, 2020). This ability forms the basis for communication and social interactions. Understanding the meaning of a multi-word utterance requires a process we refer to here as semantic binding. Lexical retrieval of information (including semantic, syntactic, and phonological details) from long term memory is required. The lexically retrieved information about single words needs to be integrated into a representation of a multi-word utterance. This process has also been referred to as merge (Chomsky, 1995; Zaccarella & Friederici, 2015) or unification (Hagoort, 2005). ERP research has demonstrated that word meaning is assembled into larger meaning representations in less than 500ms (Kutas & Hillyard, 1980), with this process immediately taking into account information from a wide range of sources, including world knowledge and discourse (Hagoort et al., 2009).

Combining words together also requires the consideration of syntactic information, including tense, aspect and agreement (Segaert, et al., 2018). Therefore, semantic binding cannot exist in the absence of syntactic binding and disentangling the two is difficult. Previous literature has disagreed on the best solution to unravel semantic from syntactic binding and therefore assigning semantic/syntactic binding-specific processes to observable effects has been difficult (Bemis & Pylkkänen, 2011). The present study examines the linguistic composition involved in adjective-noun minimal phrases. Although we refer to the observed effects as semantic binding effects, it is important to note that semantic and syntactic binding are conflated within the phrases and both processes are somewhat either simultaneously present or not.

Neuroimaging studies employing fMRI have been able to provide a wealth of information about the location of brain areas likely associated with semantic binding in young adults. Previous investigations have found evidence that semantic binding requires the exchange and integration of information in a large network of frontal and posterior areas, including left inferior frontal gyrus, bilateral superior and middle temporal gyri, anterior temporal lobe and angular gyri (Baggio & Hagoort, 2011; Lyu et al., 2019; Menenti et al., 2011; Pylkkanen, 2019; Tyler & Marslen-Wilson, 2008). The functional neural characteristics supporting specific language functions in healthy older adults, have often been found to differ from those in young adults (Antonenko et al., 2013; Peelle, 2019; Shafto & Tyler, 2014; Tyler et al., 2010; Wingfield & Grossman, 2006). With age, structural changes occur in language-relevant brain regions. In the context of these structural changes, it would be unlikely that successful performance in older adults is achieved with identical neural processes as in young adults (Peelle, 2019). Generally, the literature shows a more widespread pattern of activity in healthy older adults relative to young adults (e.g., Cabeza et al., 2002; Davis et al., 2008). Different views exist on how to interpret these age-related changes in brain activity: the appearance of more diffuse activity in older adults may reflect a general decline in neural efficiency (i.e. dedifferentiation), alternatively (though not mutually exclusive) increased engagement of brain regions may reflect focused recruitment as a means to compensate for neurocognitive decline (i.e. compensation) (Wingfield & Grossman, 2006).

One limitation of fMRI is the slow time course of the hemodynamic response (1.5-5 seconds) which limits what information it can provide about ‘when’ the specific neural processes involved in semantic processing are occurring. While EEG as a neuroimaging tool does not have the spatial resolution of fMRI, it does provide a real-time window into the neural activity underlying cognition. Previous EEG studies investigating how the brain supports semantic comprehension, have primarily looked at event-related brain potentials (ERPs) which represent brain activity phase-locked to the onset of words. As briefly mentioned above, these studies have consistently found that word meaning in young adults is integrated into the meaning of a larger multi-word utterance at around 400-500ms after the relevant word, as indexed by the N400 ERP (Kutas, Hillyard, 1980; Kutas & Federmeier, 2011). Previous studies have also elucidated several relevant aspects of how older adults comprehend sentence-level meaning. Healthy older adults do extract and make use of contextual semantic information (Stine-Morrow et al., 1999), but there are differences (compared to young adults) with respect to when and how this happens. Sentential context manipulations (i.e. the strength of contextual constraint for sentence-final words) elicit reduced and delayed N400 effects for older (compared to young) adults (Federmeier & Kutas, 2005; Wlotko & Federmeier, 2012). Moreover, effects of message-level congruity on the N400 are delayed by over 200 ms in older adults (Federmeier et al., 2003). Ageing furthermore affects processing of compositional concreteness, i.e. processing of the second noun in a noun-noun pair, in function of whether the first was concrete versus abstract (e.g. alias-battle vs. skate-battle) (Lucas et al., 2019), further suggesting that there are age-related changes in compositional semantics in healthy older (compared to young) adults.

There may be multiple (not mutually exclusive) sources of the observed age-related changes in how the brain supports making use of contextual semantic information. Older adults may engage different functional neural processes to support semantic binding and maintain a message-level meaning representation while processing incoming information. In addition, older adults may be less able to use prediction mechanisms during language comprehension. Several studies have provided support for the latter (Federmeier et al., 2002; Wlotko et al., 2012). In the present study, we focus on the former: do healthy older and young adults engage different neural mechanisms for semantic binding?

We will answer this question by investigating oscillatory activity. The EEG signal contains oscillatory activity (i.e., rhythms) which are hypothesized to play a vital role in how the brain carries out cognition (Mazaheri, et al., 2018; Siegel et al., 2012). Investigating the oscillatory (i.e., spectral) changes in the EEG allows for capturing activity that is time-locked but not necessarily phase-locked to experimental events (i.e., the onset of words). Studies focusing on the spectral changes in the EEG have found that the exchange and integration of information required for semantic binding, involves modulations in the oscillatory power in the theta (~4-7 Hz), alpha (~8-12 Hz), and beta (~15-30 Hz) bands (for comprehensive reviews: Meyer, 2018; Prystauka & Lewis, 2019; for a detailed overview see Weiss & Mueller, 2012). Modulations of each frequency band are thought to reflect different language comprehension related processes. However, it is not always easy to map specific roles onto definitive oscillatory ranges. The theta frequency range is related to memory retrieval and processing demands, whereas alpha parallels attentional processes and storage of phrases (Meyer, 2018). Previous studies have also further subdivided the beta frequency ranges into more narrow bands: ~13-20Hz (i.e., low beta), and 20-30Hz (i.e., high beta) (Poulisse et al., 2020; Segaert, Mazaheri, et al., 2018; see Weiss & Mueller, 2012 for a detailed overview). Low beta is related to higher-order processing. In language comprehension this translates to linking past and present input (i.e. binding including semantic features (Bastiaansen & Hagoort, 2006; von Stein et al., 1999; Weiss & Mueller, 2003) and syntactic unification (Bastiaansen et al., 2010)). High beta has previously been related to processing of action/motor-related language (Elk et al., 2010).

Although there are a number of previous studies that reveal the oscillatory signatures of semantic binding in young adults, the pattern across studies is not always clear cut. Firstly, several previous studies (but not all, e.g., Kielar et al., 2018; Wang et al., 2018) have shown that increased theta power is associated with semantic anomalies (Bastiaansen & Hagoort, 2015; Davidson & Indefrey, 2007; Hagoort et al., 2004; Hald et al., 2006; Wang, Zhu, et al., 2012). This is thought to reflect the increased effort (i.e. neural resources) required to integrate semantically incongruous items into the wider context. Furthermore, increased theta power has also been linked with syntactic violations (e.g., Bastiaansen et al., 2002; Kielar et al., 2015; Lewis et al., 2016), supporting the notion that this theta signature may reflect a more general violation detection mechanism (Prystauka & Lewis, 2019). Furthermore, the importance of beta oscillations in linguistic composition processing has previously been highlighted (Lewis et al., 2015; Lewis & Bastiaansen, 2015). Lewis and Bastiaansen (2015) proposed the hypothesis that power changes in the beta band are linked to linguistic information maintenance. More specifically, active maintenance of linguistic cues leads to beta power increases. The language processing system actively maintains the current cognitive state due to the greater processing demands arising from more complex clauses. This notion is supported by a study by Meyer et al. (2013), who have found an increase in the beta power (13 to 20Hz) at the point of the verb in sentences with long-distance dependencies (between argument and verb) compared to sentences that contained short-distance dependencies. Long-distance dependency clauses are computationally more demanding and complex (compared to short-distance sentences), directing the language comprehension system towards active maintenance of the processing mode.

On the other hand, any change of linguistic input is related to beta power decreases. Lewis and Bastiaansen (2015) suggested that semantic (and syntactic) violations are clear cues to the system indicating a need for change. Therefore linguistic violations lead to decreases in beta power compared to instances where semantic (or syntactic) violations are not present. This theory is supported by multiple empirical studies. For example Kielar et al. (2014) (see also Kielar et al., 2015, 2018) investigated the effect of violations on oscillatory responses using semantically correct (e.g. “A new computer will *last* for many years”) vs. semantically incorrect (e.g. “A new computer will *paint* for many years”) sentences. They found that the semantic violations elicited power decreases in the 8 to 30Hz range and were maximal ~500-1000 ms post the target word onset over parietal sites. Furthermore, Luo et al. (2010) provided additional support for the notion that semantic violations lead to low beta (16 to 20Hz) decreases immediately after (0-200 ms) the target word onset as well as in a later window (~500 ms later) using Chinese semantically congruent vs. incongruent sentences. This beta modulation in the later time window (around the N400 effect) has also been found by Wang, Jensen et al. (2012) with anomalous words eliciting a decrease in the beta power. On the other hand, Lam et al. (2016) reported an opposite result, showing stronger beta power decrease for real sentences compared to word lists with the most prominent effect at ~350ms. The authors related this effect to stronger neural activation for sentences (compared to word lists) reflecting the unification of semantics and syntax. Lastly, several previous studies suggested that the beta signature for maintenance and binding of linguistic information extends into the alpha power range (Gastaldon et al., 2020; Kielar et al., 2014; Lam et al., 2016; Luo et al., 2010; Segaert, Mazaheri, et al., 2018).

In the present study, we aim to investigate modulations in oscillatory brain activity (with pre-defined frequency bands based on previous literature: theta (4-7Hz), alpha (8-14Hz), low beta (15-20Hz), and high beta (20-25Hz)) during semantic binding in healthy older compared to young adults, using a minimal two-word phrase paradigm. We present target words (e.g. horse) in a semantic binding (e.g. swift horse) vs. no semantic binding context (e.g. swrfeq horse). In both cases, for the target word, retrieval of lexico-semantic information from memory takes place. However, only in the binding condition a complex meaning representation can be built for the phrase, based on the elementary building blocks of each individual word. It is important to note here again that syntactic binding is also present in the semantic binding context as the two are not easily disentangled. As the semantic content, and not the syntactic features of the phrases, is manipulated in the present paradigm, we refer to the conditions as semantic versus no semantic binding. Within the semantic binding condition we also manipulate whether the phrase is plausible. Although secondary, the use of this paradigm also allows an investigation of lexical retrieval effects and the recognition of the word form (e.g. swift vs. swrfeq) so we will report these findings as well.

The computation for a two-word phrase forms the foundation of binding in the context of increasing complexity. Investigating elementary semantic binding by means of a minimal phrase paradigm offers the advantage of focusing on the binding process while minimizing contributions of other processes involved in sentence comprehension, such as working memory load and the ability to use predictions. This advantage is particularly salient when investigating age-related changes in how the brain supports online sentence comprehension, given that working memory and the ability to use predictions are also impacted by age. Bemis & Pylkkänen (2011) conducted one of the first studies with a minimal paradigm, and compared nouns in a minimal binding context (e.g. red boat) versus a wordlist condition (e.g. cup boat). This inspired many other studies to use similar designs (Bemis & Pylkkänen, 2013; Pylkkänen et al., 2014; Segaert, et al., 2018; Zaccarella et al., 2017; Zaccarella & Friederici, 2015). Poulisse et al., (2019, 2020) used this approach to investigate how healthy ageing impacts minimal syntactic binding.

In line with previous literature on semantic comprehension in young adults (Lewis et al., 2015), we expected to see a greater beta power decrease (particularly in the low-beta (15-20Hz) frequency band) in the no semantic binding compared to the semantic binding condition. We made no predictions concerning a modulation in the theta range, since our paradigm does not manipulate violations (Prystauka & Lewis, 2019), but rather, successful binding versus no binding (with previous behavioural performance results demonstrating that no binding occurs for pairing a pseudoword with a real word (Poulisse et al., 2019; Segaert, Mazaheri, et al., 2018)). If previously observed age-related changes in making use of contextual semantic information are at least in part due to a change in how the brain supports semantic binding, then we expect to see different oscillatory signatures for healthy older versus young adults, in the semantic binding versus no semantic binding conditions, however this aspect of our study is exploratory and entirely novel, making it difficult to make concrete predictions about the direction of power changes in the alpha or beta range (Beese et al., 2019; Poulisse et al., 2020).

## Materials and methods

### Participants

33 young adults and 32 healthy older adults took part in the study. However, 7 participants were excluded from the analysis due to: (a) excessive EEG artefacts in the recordings (N=4), and (b) being bilingual (N=3). The participants included in the analyses were 29 young adults (2 males, aged 18-24) and 29 healthy older adults (13 males, aged 63-84) (see Table 1 for more information). All participants were right-handed, British-English monolingual speakers with normal-to-corrected vision and no neurological or language impairments. All older adults scored above 26 out of 30 in the Montreal Cognitive Assessment (MOCA) test (M = 27.79, SD = 1.01) (scores ≤ 26 suggest risk of mild cognitive impairment or dementia (Smith et al., 2007)).

**Table 1.** Demographic and cognitive characteristics for young and older adult participants

|  | Young adults<br>(N=29) |  | Older adults<br>(N=29) |  | *t-value* | *P* |
| --- | --- | --- | --- | --- | --- | --- |
|  | *M* | *SD* | *M* | *SD* |  |  |
| Age (years) | 19.5 | 1.5 | 73.6 | 5.8 |  |  |
| Years of education | 14.52 | 1.5 | 15.45 | 3.08 | -1.46 | .15 |
| Processing speed | 81.48 | 11.57 | 65 | 13.17 | 4.97 | < .001 |
| NART | 25.81 | 5.07 | 37.05 | 4.61 | -8.82 | < .001 |
| Working memory | 4.42 | .66 | 4.97 | .92 | -2.62 | < .05 |

The young adults were Undergraduate students from the University of Birmingham and took part in the study for course credits. The older adults were from the Patient and Lifespan Cognition Database and were compensated for their time with cash payments. Participants signed informed consent, which followed the guidelines of the British Psychology Society code of ethics, and the experiment was approved by the Science, Technology, Engineering, and Mathematics (STEM) Ethical Review Committee for the University of Birmingham (Ethics Approval Number: ERN_15-0866).

There was no significant difference in the number of years spent in education between the younger and the older adults. In line with expectations, young adults outperformed older adults in processing speed (Weschler Adult Intelligence Scale-IV processing speed index), whereas older adults outperformed young adults on the National Adult Reading Test (Nelson, 1982). Surprisingly, older adults also outperformed young adults in the working memory tasks (i.e. the average combined score of the backward digit span and subtract 2 span tests) (Waters & Caplan, 2003), which could be attributed to young adults being less motivated when they were participating in the tasks (in line with similar findings reported previously: Heyselaar et al., (2020)).

### Design, materials and task

We created a minimal language comprehension paradigm with two-word-phrases. Each phrase included two words, where the target word was always the second word. The design of the study, with example stimuli, is illustrated in Figure 1. We manipulated lexical retrieval (comparing real words to letter strings) and semantic binding (comparing the target word in a semantic binding context to a no semantic binding context). We were primarily interested in the effects of semantic binding, but since our two-word phrase paradigm allows examining the effects of lexical retrieval and the recognition of the word form also, we report these effects below as well. Within the semantic binding condition, we furthermore manipulated whether semantic binding was plausible (e.g. swift horse) or implausible (e.g. barking horse).

**Figure 1.**
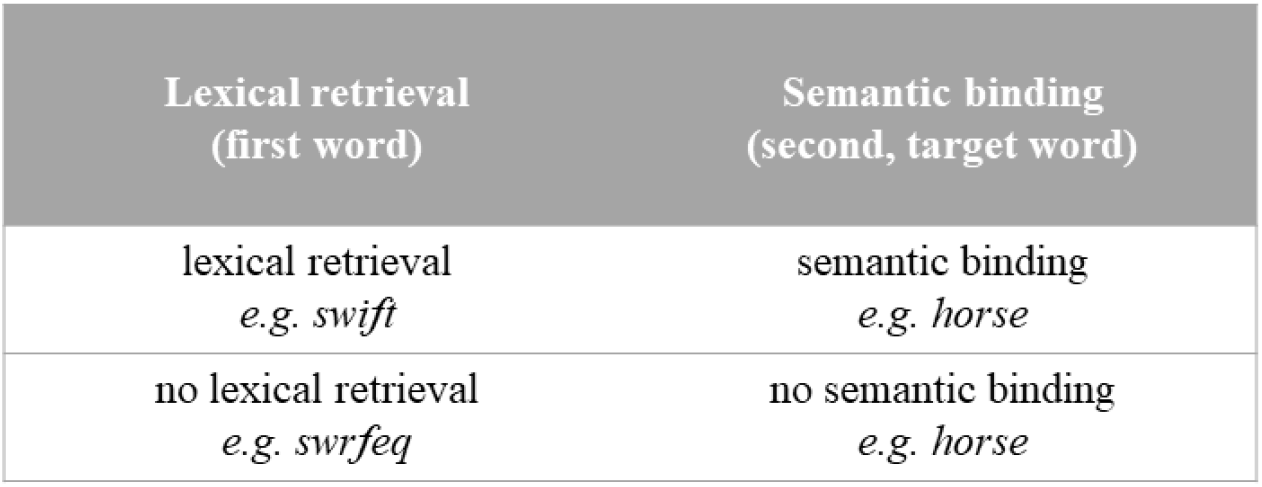
Example word-pairs in each condition.

To ensure participants paid attention to the word-pair stimuli throughout the experiment, we included questions about the word-pairs on a subset of the trials (22% of all trials). The questions asked “Did you just see [word pair]”. There were no significant differences between young and older adults in response accuracy (young adults: mean = 94.83, SD = 0.5; older adults: mean = 96.48, SD = 0.2; t(42.109) = −1.557, p = .127) or reaction times (young adults: mean = 1718.78, SD = 503.64; older adults: mean = 1851.45, SD= 488.92; t(56) = −1.018, p = .313). All the participants scored higher than 80%. From this we can conclude that young and older adults paid close attention to the language stimuli as they were being presented to them throughout the experiment.

We verified our plausibility manipulation in an online rating study with 57 respondents. The online survey asked to rate the plausibility of the two-word phrases, where 1 = ‘Completely implausible’, 2 = ‘Somewhat implausible’, 3 = ‘Somewhere in between’, 4 = ‘Somewhat plausible’, and 5 = ‘Completely plausible’. The plausibility ratings were significantly different for the plausible (M=3.87, SD=.29) and the implausible word-phrases (M=1.93, SD=.66); Welch’s F(1, 101.65) = 526.36, p < .001.

In each condition, about half of the target words were animate, the other target words were inanimate. The exact trial distribution was as follows: inanimate-plausible (N=45), inanimate-implausible (N=46), inanimate-letter string (N=44), animate-plausible (N=42), animate-implausible (N=47), and animate-letter string (N=46). The list of plausible and implausible adjectives was matched for word frequency using the CELEX database (Baayen et al., 1993) (plausible *mean* = 28.16, *SD* = 40.63, implausible *mean* = 27.87, *SD* = 36.67), number of syllables (plausible *mean* = 1.74, *SD* = 0.49, implausible *mean* = 1.79, *SD* = 0.52) and number of letters (plausible *mean* = 5.87, *SD* = 0.79, implausible *mean* = 5.64, *SD* = 0.71).

Three versions of the experiment were created, where the same word-pairs were presented in different orders. Participants were randomly assigned to one of the three versions. The paradigm intended to present 60 attention-questions for each of the versions of the experiment. However, due to an error in creating the question lists, the number of questions differed slightly per version (either 61 or 62 questions). The questions were not used in any of the EEG analyses. A full stimulus list, each of the 3 versions of the experiment, with the exact attention questions asked, can be downloaded from https://osf.io/f8grv/.

### Procedure and trial timing

We presented our experiment using E-prime 2.0. Figure 2 depicts the duration of time that each element of the trial was presented for (top row) and the presentation time of each element in the trial when the EEG epoch was locked to the onset of word 1 (bottom row). The task consisted of 270 trials divided into 9 blocks. In between each block, we offered the participants a break.

**Figure 2.**
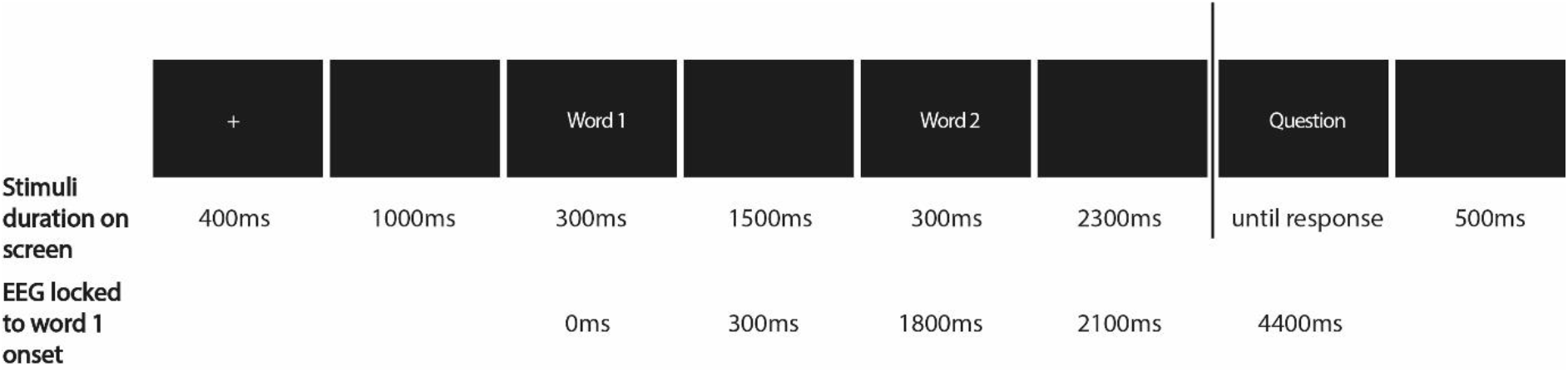
Trial presentation of the minimal two-word phrase paradigm. The questions appeared in 22% of the trials. The top row (“Stimuli duration on screen”) depicts the on-screen time duration (in ms) of each trial element. The bottom row (“EEG locked to word 1 onset”) depicts the presentation time (in ms) of each trial element when the trial/ EEG epoch is locked to the onset of word 1.

Upon the start of the experimental session, participants were fitted with a 64-electrode EEG cap. Once the EEG set-up was finished participants sat in a sound proof booth 70 cm from the monitor where the computerised task took place. Participants were instructed to read in silence word-pairs (e.g., swift horse) appearing on the screen. They were told that from time-to-time they would see a question on the screen regarding the word-pair that they had just seen (e.g. Did you just read ‘swift horse?’). Participants were able to indicate ‘yes’ or ‘no’ using a button box. Participants completed a practice block first to familiarise themselves with the paradigm (30 word-pairs, with 9 questions, which were different from the experimental stimuli), which was followed by the actual experiment. Following the computer task, participants completed Working Memory tests (i.e. the Backward digit span task and the subtract 2-digit span task) (Waters & Caplan, 2003), National Adult Reading Test (NART) (Nelson, 1982), and the Weschler Adult Intelligence Scale-IV processing speed index (Weschler, 2008). In addition, the older participants also completed the Montreal Cognitive Assessment (MoCA) (Nasreddine et al., 2005).

### EEG recording

EEG was recorded using Waveguard caps containing 64 cap-mounted Ag/AgCI electrodes (10-20 layout, including left and right mastoids). Horizontal eye movements were measured by two electrodes placed on the outer left and right canthi. Vertical eye movements were recorded by two electrodes placed above and below right eye. The EEG recording was acquired with online reference to the CPz channel. The signal was amplified with the ANTneuro EEGosports amplifier system and recorded using EEGo software (Advanced Neuro Technology). The signal was obtained at a sampling rate of 500Hz, with a 30Hz low-pass filter (24 dB/octave) and a 0.05Hz high-pass filter, implemented in the EEGosports firmware. We aimed to keep the impedances below 10 kΩ.

### EEG analysis

The EEG pre-processing was performed using EEGLAB 14.1.2b (Delorme & Makeig, 2004) and Fieldtrip toolbox 2018-07-16 (Oostenveld et al., 2011). The data were epoched to the onset of the first word (−1.5sec to 4.4sec) and later offline re-referenced to the average of all of the channels, where the mastoid and bipolar electrodes were excluded from the re-referencing. EEGLAB was used for manual inspection and rejection of trials with non-physiological artefacts. The average number of removed trials was 26.34 (SD=21.93) per participant due to artifacts. Ocular artifacts were removed based on the scalp distribution using independent component analysis (“runica”) in EEGLAB. The average number of removed components was 2.07 (SD=.81) for each participant. Channels TP8 and TP7 were removed before completing any analyses due to poor or no signal from these channels across the participants.

### Time-frequency representations of power

Time-frequency representations (TFRs) of power were performed using the FieldTrip ‘mtmconvol’ method with a sliding time window. The Hanning taper was applied to the adaptive time window of 3 cycles per each frequency of interest (i.e. the length of the window at each frequency of interest is equal to 3/f s) for every trial. Similar approaches were used previously by Mazaheri et al., (2009); Poulisse et al., (2020); van Diepen et al., (2015). The analysis included the frequency of interest of 2Hz to 30Hz in steps of 1Hz, and the time of interest of −1.5 to 4 sec in steps of .05 sec. We calculated the changes in oscillatory power locked to the onset of the stimulus (i.e. word one) in relation to the change in power from baseline. The data were baseline corrected to a window of −600 to −100 ms prior to stimulus onset (i.e. presentation of first word). This was predefined and thus applied within conditions and age groups.

We statistically examined baseline differences between groups (collapsed across conditions) using non-parametric cluster-based permutation tests (see below for further explanation) prior to correcting the baseline window. We used an averaged priori time window of −600 ms to 0 sec (where 0 was the onset of word 1) within the following pre-defined frequency bands: theta (4-7Hz), alpha (8-14Hz), low beta (15-20Hz), and high beta (20-25Hz) in the cluster-based permutation tests. The cluster-based permutation tests did not reveal any significant differences between groups (healthy older adults vs. young adults collapsed across all conditions) in the averaged baseline window (−600 ms to 0).

To ensure that the observed oscillatory changes were not just the spectral representation of the ERPs, the ERP components were subtracted from the TFR (Mazaheri & Picton, 2005). The subtraction was achieved by first generating the time frequency decomposition of the ERP data for each condition and participant separately. Next, the time frequency power spectra of the ERP was subtracted from the time frequency power spectra of the EEG signal for each condition. The subsequent power changes in the time-frequency domain were used to generate time frequency power spectra differences between experimental conditions (lexical retrieval/semantic binding vs no lexical retrieval/ semantic binding; plausible semantic binding vs implausible semantic binding) for each group separately.

Finally, the statistical differences of the experimental condition differences in the power changes in the time-frequency domain were assessed by using a non-parametric cluster-based permutation test (using FieldTrip toolbox) (Maris & Oostenveld, 2007). Each channel/time/frequency pair locked to the onset of the first word for the difference between each experimental condition (i.e. binding vs no binding) was compared using an independent (two-tailed) samples t-test (for young vs older adults) with a threshold at 5% significance level. Significant pairs were then clustered (cluster was defined based on proximity in space using the triangulation method i.e., having at least two significant electrodes that were adjacent to each other) and participants’ labels of each cluster were randomly shuffled using 1000 partitions. The Monte Carlo P values were calculated using the highest sum of the test statistic. An equivalent dependent (two-tailed) samples t-test was used to compare the experimental manipulations within each group separately in order to extract between condition effects. The time window used to assess the statistical differences in time-frequency power for (1) lexical and semantic binding manipulation was 0 (onset of word 1) to 3.2 sec, where onset of word 2 occurred at 1.8 sec, and (2) plausibility manipulation was 1.8 sec (onset of word 2) to 3.2 sec. The above analysis was performed within the following pre-defined frequency bands: theta (4-7Hz), alpha (8-14Hz), low beta (15-20Hz), and high beta (20-25Hz) consistent with previous studies (Poulisse et al., 2020; Segaert, et al., 2018). The average number of trials across all participants included in the final analysis for the lexical retrieval/ semantic binding condition was 162.87 (SD = 15.27), for the no lexical retrieval/ no semantic binding condition 80.79 (SD = 7.4), for the plausible condition 79 (SD = 7.6), and for the implausible condition 83.86 (SD = 8.05). Note here that we checked whether any of the observed condition effects in the data are not spurious differences attributed to the imbalance of signal-to-noise ratio (SNR) (due to the imbalance of the number of trials in lexical retrieval/semantic binding and no lexical retrieval/ no semantic binding). To do this, we used a random re-sampling approach to equate the trials in both conditions. We then completed between condition (lexical vs. non lexical) dependent non-parametric cluster-based permutation tests within each group separately and found similar condition effects in each group (see supplementary materials for the full report of these results, and Suppl. Figure 2) as those reported here. Therefore, we can be confident that any observed condition differences in each group are not due to the imbalance of SNR.

## Results

We first visually inspected the TFRs and qualitatively describe the power modulations. The onset of word 1 and word 2 generated an increase in theta (4-7Hz) power, followed by a suppression of alpha power (8-14Hz), irrespective of condition in both age groups (Figure 3). Consistent with previous studies (e.g., Bastiaansen et al., 2005, 2008; Hermes et al., 2014; Mazaheri, et al., 2018), the theta power increase peaked at around 0.2 sec post word onset and was maximal over the occipital channels. Also in line with previous work, the alpha power suppression peaked at around 0.5 sec post word onset and was maximal over the occipital channels (Davidson & Indefrey, 2007; Mazaheri, et al., 2018).

**Figure 3.**
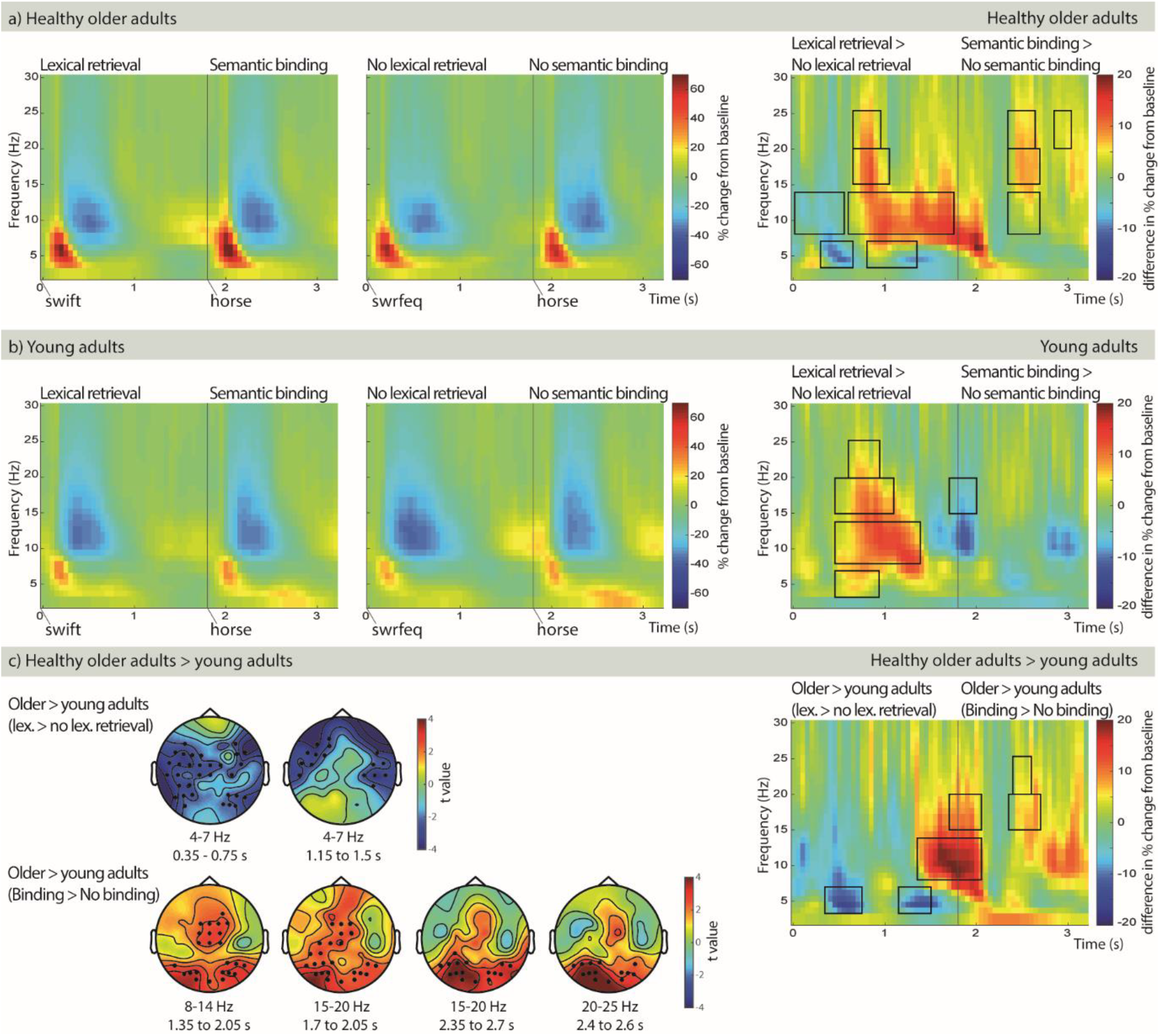
TFRs of power (collapsed across all electrodes) for lexical retrieval/semantic binding and no lexical retrieval/no binding, in (A) the healthy older adults, and (B) the young adults. (C) The condition differences (i.e. lexical retrieval/ semantic binding – no lexical retrieval/ no semantic binding) in older adults minus the condition differences in young adults. Head plots are illustrating the clusters of electrodes that show the most pronounced mean condition difference for the healthy older adults vs. the young adults. Black rectangles indicate significant group differences (p<0.05, cluster corrected).

Our analysis approach was as follows. We focused our analysis on the oscillatory changes in the EEG associated with lexical processing (e.g., 1^st^ word: swift vs. swrfeq) and semantic binding (e.g., horse when preceded by swift vs. when preceded by swrfeq). To help with the interpretation of the between-group effects, in the main text we only describe the significant oscillatory differences between conditions within groups (Figure 3A and B) *if* they were also significantly different between young and older adults (Figure 3C). A comprehensive and more detailed description of condition differences within each age group separately can be found in the supplementary materials (and Suppl. Figure 1).

In what follows, we first describe the lexical retrieval results, and then the semantic binding results.

### Lexical retrieval results

We investigated the effect of lexical retrieval and recognition of the word form through comparison of power differences in the word 1 time window, comparing real words (i.e. lexical retrieval) to letter strings (i.e. no lexical retrieval can successfully take place (e.g., swift vs. swrfeq)). We observed significant differences between age groups for the lexical retrieval effects, and clusters in the observed data were found in the theta (4-7Hz) and alpha (8-14Hz) range (Figure 3C). We discuss these in turn below.

#### Theta and alpha power modulations for lexical processing show opposing patterns for healthy older adults and young adults

We observed significant differences in theta activity between the age groups (p = .014) during lexical retrieval (i.e. lexical retrieval minus no lexical retrieval) maximal over occipito-temporal electrodes. Specifically, we observed that theta was attenuated around 0.35 to 0.75 sec after the first word during lexical retrieval in the healthy older adult group relative to the young adult group. This age-group difference in theta activity during lexical retrieval emerged due to opposing patterns of theta modulation in relation to lexical and no lexical processing trials. Specifically, within the healthy older adults there was a smaller increase in theta power (p = .02) for lexical compared to no lexical retrieval in a corresponding time window, maximal over right occipital and left central electrodes (Figure 3A). However, in contrast the young adults showed a greater increase in theta power for lexical compared to no lexical retrieval (p = .02) in a similar time window (0.45 to 0.95 sec), maximal over left occipital and parietal electrodes (Figure 3B).

In addition, we observed differences in theta power related to lexical processing (p = .05) corresponding to a cluster extending between 1.15 to 1.50 sec, with the power being attenuated in the older adult group compared to the young participants. This age-group difference, pronounced over frontal-lateral electrodes, emerged due to an increase in theta activity in the non-lexical relative to the lexical condition (p = .004) in a cluster in the observed data that extended from 0.8 to 1.35sec in the elderly participants, which was absent in the young group.

Last, we observed a significant group difference (p = .034) in the alpha modulation (8-14Hz) that corresponded to a cluster extending from 1.35 to 2.05 sec. This observed cluster was maximal over occipital and central electrodes. This group difference emerged due to an opposite pattern of alpha power rebound during lexical processing between the older and young adults. In the older adults the alpha power rebound following the post-word alpha suppression was greater in the lexical retrieval trials than the no lexical retrieval trials (p = .002). A cluster in the observed data spanned from 0.6 to 1.75 sec. On the other hand a reversed pattern was seen in the young participant group with the alpha rebound being greater in the non-lexical retrieval trials compared to the lexical retrieval trials (note here that this is a qualitative description of the power modulations based on visual inspection only, rather than a significant effect). We interpret the attenuated alpha rebound to reflect an absence of closure in the no lexical retrieval condition in the older adults. Specifically, we hypothesize that the older adults continue to try and retrieve a lexical item (for a longer time than the young adults) after the onset of a pseudo word. Finally, it must be noted that the alpha rebound overlaps with the onset of the second word (i.e. the word to be semantically integrated). It is possible therefore that the between-group difference in alpha rebound must be partly attributed to semantic binding (discussed further below) rather than exclusively lexical processes.

#### Lexical retrieval results summary

Summarizing above significant between-group differences in lexical retrieval and recognition of the word form, we find that the older and young adults appear to exhibit opposite patterns of theta and alpha modulation after the onset of real words (i.e. lexical retrieval) vs letter strings (i.e. no successful lexical retrieval). Specifically, closely followed by word onset, older adults exhibited a smaller, whereas young adults exhibited a larger theta power increase, for the lexical compared to the non-lexical condition. Additionally, the alpha power rebound effect was reversed between the groups: in the older adults it was greater, and in the young adults smaller (though not significantly so), in the lexical retrieval condition compared to the non-lexical condition.

### Semantic binding results

Semantic binding effects were defined by comparing oscillatory power surrounding the onset of a second (i.e. target) word in a semantic binding to a no semantic binding context (e.g., *horse* when preceded by *swift* vs. when preceded by *swrfeq*). We observed significant differences between age groups for the semantic binding effects, with clusters in the observed data being found in the alpha (8-14Hz), low beta (15-20Hz) and high beta range (20-25Hz) (Figure 3C). We discuss these in turn below.

#### Alpha and beta power in a time window preceding and following the onset of the target word was differentially modulated across the age groups

A cluster in the observed data was found in the alpha band (8-14Hz), which corresponded to a significant between-group difference (p = .034). The observed cluster extended from 1.35 to 2.05 sec over occipital and central electrodes. Although within-groups there were no significant condition differences, for young adults we observed that the alpha rebound was attenuated preceding and following the target word in the semantic binding condition compared to the no semantic binding condition. On the other hand for older adults in the equivalent time window an opposite pattern was observed with the alpha rebound being larger in the semantic binding relative to the no semantic binding trials. The pattern just described in the alpha range, extends into the lower beta range (15-20Hz). There was a significant between-group effect (p = .046) with a cluster extending from 1.7 to 2.05 sec, maximal over occipital, parietal and central electrodes. For young adults, there was a smaller low-beta increase in the semantic binding compared to the no semantic binding condition (p = .036), corresponding to a cluster that extended from 1.7 to 2sec.

#### A beta rebound was observed for semantic binding for the healthy older adults but not for the young adult group

In addition, there were between-group effects that mapped onto clusters found in the lower beta (15-20Hz) and higher beta ranges (25-30Hz) over occipital electrodes that extended from 2.35 to 2.7 sec and 2.4 to 2.6 sec respectively. The between-group effect in the lower-beta (p= .002) and higher-beta range (p = .046) is driven by the older adults showing a clear semantic binding signature in this time window (with the effect extending into the alpha range), with no effect for the young adults in the equivalent time window. The healthy older adults elicited greater and more sustained suppression (p = .002) of lower-beta, and more higher-beta suppression (p = .002) in the no semantic binding condition compared to the semantic binding condition. These effects corresponded to clusters that extended from 2.35 sec and ended around 2.65 to 2.7 sec.

#### Semantic binding results summary

Summarizing above significant between-group differences in semantic binding, we see that young and healthy older adults clearly have different semantic binding signatures. In young adults, there is an attenuation of the alpha rebound in anticipation of the target word onset, followed by a binding signature in the low-beta band (i.e. a smaller low-beta increase in the semantic binding compared to the no semantic binding condition) immediately preceding and during the presentation of the target, to-be-integrated, word. In contrast, during the semantic binding condition the older adults exhibited a smaller decrease in high and low beta activity (compared to no semantic binding) starting only at 500 ms after the onset of the target word, which was not present in the young adults.

### No effects of plausibility

We carried out a non-parametric cluster-based permutation analysis within the 1.8 to 3.2 sec time window of interest with pre-defined frequency bands (described above) to compare plausible semantic binding (e.g. swift horse) to implausible semantic binding (e.g. barking horse). We found no significant differences in the power changes in any of the pre-defined frequency bands between healthy older and young adults. Important is that we do not take this to suggest that there are no age-related differences between young and older adults in the effects of plausibility on semantic integration, but rather, that our experiment was potentially not suited to reveal them. We must note that within the older adults group, the condition comparison between plausible and implausible semantic binding elicited no power differences; within the young adults a power difference approached significance (p = .054), corresponding to a cluster that extended only from 2 to 2.15 sec. This observed cluster was found in the low beta (15 to 20Hz) band. It is likely therefore that our plausibility manipulation was not strong enough to elicit reliable condition differences, in either age-group, and therefore our experimental manipulation may not have been sensitive enough to investigate potential age-related changes in processing plausibility.

## Discussion

The current study used a minimal two-word phrase paradigm to investigate the differences in oscillatory activity (in the theta, alpha, and beta range) during lexical retrieval and semantic binding in healthy older vs. young adults. Lexical retrieval was assessed by comparing neural patterns during the presentation of real words (e.g., swift) vs. letter strings (e.g., swrfeq). Semantic binding was examined by comparing neural patterns between semantic binding (e.g., horse, preceded by swift) and no semantic binding (e.g., horse, preceded by swrfeq) conditions. Here, it is important to highlight that although the present study manipulated the semantic binding context of the phrases, syntactic binding was also present in the semantic binding condition. Therefore, the semantic binding effects likely cover both semantic and syntactic compositional properties.

With regards to lexical processing we found that the older and younger groups exhibited opposite patterns of theta and alpha modulation at specific time intervals after word onset, which as a combined picture suggest that lexical retrieval is associated with different and delayed signatures in older compared to young adults. Interestingly, with respect to semantic binding, we observed a signature in the low-beta range for young adults (i.e. a smaller increase for semantic binding relative to no binding) surrounding the presentation of the target word, while the semantic binding signature for older adults occurred about ~500ms later as a smaller low- and high-beta decrease (for binding compared to no binding). We will now discuss each of these findings in more detail in relation to previous literature.

### Age-related oscillatory patterns linked to lexical retrieval

Firstly, we found that the oscillatory patterns observed during lexical retrieval (i.e. post word one onset) were different for healthy older adults compared to young adults. The presentation of word one (regardless of condition) led to an increase in theta power across both age groups, which has previously been proposed to be linked with the role of long-term memory retrieval (Bastiaansen et al., 2002, 2008; Bastiaansen & Hagoort, 2006).

The lexical retrieval effect (i.e. a real word vs letter string) was associated with a greater theta increase closely followed after the first word onset (0.45 to 0.95 sec) in young adults over the left temporal and parietal sites, suggesting greater demand on retrieving the meaning of ‘real’ words from long term memory. In other words, the effort required to retrieve an item containing lexical information was greater compared to when the item lacked lexical representation. This is consistent with previous studies (Bastiaansen et al., 2005; Marinkovic et al., 2012; Mellem et al., 2013) who found that items that carry greater meaning or are more complex (i.e. real words and open class words) elicit stronger theta response compared to items that lack (or have lesser) lexical representation (i.e. pseudo words and closed class words) in young adults. In Shahin et al., (2009) participants made voice identification or semantic judgements to auditory word stimuli and found that theta power is heightened for the latter. Although the current study is visual, it appears that a similar mechanism is observed as for the auditory modality.

In contrast, amongst healthy older adults, the lexical retrieval effect was associated with a theta increase which was smaller closely after word one onset (0.3 to 0.65 sec) and then bigger (0.8 to 1.35 sec) in the lexical retrieval condition (vs. no lexical retrieval). In other words, the ‘typical’ lexical effect (i.e. greater theta increase in the lexical retrieval compared to no lexical retrieval condition) occurred ~350ms later in healthy older adults compared to young adults and was more widely spread. We suggest that healthy older adults require a longer time and a wider network to retrieve a lexical item from memory compared to young adults. However, it should be noted that this is a highly speculative conclusion as the topography of this lexical retrieval effect is varied between the age groups.

The lexical retrieval effect in healthy older adults in the theta band is contrary to Mellem et al., (2012) who did not find any theta power differences associated with lexico-semantic processing when comparing open and closed class words in older adults (although the lexical manipulations differ between the current study and Mellem et al., (2012), both paradigms manipulated the level of lexico-semantic content). Additionally, this theta power difference between lexical vs. no lexical retrieval was prominent over a more widely spread network in healthy older adults including the bilateral occipital and central electrodes compared to young adults where it was lateralised to the left temporal and parietal sites. This is in line with the frequently found tendency for healthy older adults to show a lesser engagement of task relevant regions but a greater involvement of other regions compared to young adults (Cabeza et al., 1997, 2002; Grady, 2000). Furthermore, the Hemispheric Asymmetry Reduction in older adults (HAROLD) model (Cabeza, 2002) suggested that the neural processing in healthy older adults is associated with a decrease in hemispheric asymmetry (whereas young adults show lateralization to one side), which is evident in the theta activity in the current study.

It should be noted here that linguistic processes other than lexico-semantics are involved in reading words vs. letter strings, including recognition of the word form and orthographic processing (Taylor et al., 2013); thus the results relating to the manipulation of lexical retrieval will also refer to these features. The left ventral occipito-temporal (vOT) cortex has previously been linked to word processing, specifically to aligning to orthographic features (Schurz et al., 2014). Froehlich et al. (2018) found that age-related differences in word processing were most pronounced during orthographic processing in (among others) the vOT circuit. This is consistent with our present age-related theta effect as this was most pronounced at the occipito-temporal sites. This suggests that the age-related difference in processing words vs. letter strings speculatively lies in the recognition of the word form and its orthographic properties.

Lastly, older adults did not show an alpha rebound like the young adults: while young adults showed an alpha rebound that was greater in the no-lexical retrieval compared to the lexical retrieval, the older adults showed the opposite. We interpret this to reflect an absence of closure in the no lexical retrieval condition in the older adults: older adults may continue to try and retrieve a lexical item (for a longer time than the young adults) after the onset of a pseudo word. In other words, processing of the non-lexical item was not fully complete, consistent with a finding in previous ageing studies using a paradigm with pseudo words (Poulisse et al., 2020). However, this interpretation is tentative and we elaborate on this in the next section.

### Age related oscillatory patterns linked to semantic binding

Most interestingly, we observed oscillatory differences between healthy older and young adults during and after word two onset. The target word (i.e. the second word) in the semantic binding condition required participants to retrieve the lexico-semantic information from memory, and, to develop (i.e. bind together) a compound meaning representation of the two-word phrase. The latter was absent in the no semantic binding condition where the combination of a letter string and a real word cannot create a meaningful phrase. We observed clearly different semantic binding signatures (i.e. comparing the target words in the semantic binding vs. no semantic binding condition) for the young vs. healthy older adult groups. Although we refer to the below effects as semantic binding signatures, it is important to note here again that the comparisons of the respective conditions either simultaneously contained syntactic and semantic binding or no presence of binding at all. These are novel findings as few previous studies have investigated the effect of ageing on the oscillatory dynamics associated with semantic binding.

The semantic binding effect was associated with an oscillatory brain activity difference in the alpha (1.35 to 2.05 sec) and low-beta (1.7 to 2 sec) frequency bands between healthy older and young participants. The effects that occur around the onset of the second word must be treated with caution. Due to the caveats of the experimental design, one cannot easily map these effects onto either the lexical status of the first word or the anticipatory processing related to binding of the second word. We can only present plausible explanations for the observed results based on the comparisons to previous literature. Therefore, here we speculatively propose that the first part of the group difference in the alpha band (up until around 1.7 sec) occurred due to older adults eliciting an atypical alpha rebound response in the no lexical retrieval condition, whereas the young adults had no condition effect in the same time window (already discussed in the previous section on lexical retrieval). The later part of this alpha effect (i.e., post second word onset) may potentially be implicated in the binding process itself, including making semantic predictions. Although previous research (e.g., Luo et al., 2010) has related alpha power modulations with a violation of semantics (and therefore binding), these effects occur 400-600 ms after the onset of the target word and not during it. For this reason, we cannot definitively conclude whether this alpha power modulation surrounding the onset of the second word is related to the lexical retrieval closure of the first word, the anticipatory processing of binding, or a combination of the two.

Importantly, young adults elicited a smaller low-beta increase in the semantic binding condition (vs. no semantic binding) in a time-window immediately preceding and during the presentation of the to-be-integrated target word (from 1.7 to 2 sec for low-beta). This early binding signature in the young adults within the beta band is somewhat consistent with previous studies. Beta frequencies have previously been proposed to be “carriers” of linguistic information and are involved in binding past and present inputs (Weiss & Mueller, 2012). For example, von Stein et al., (1999) found coherence exclusively in the beta frequency range between left temporal and parietal sites during semantic binding across visual and auditory modalities. Additionally, Berghoff et al., (2005) showed that figurative compared to literal sentences elicited increased coherence in the beta band between the hemispheres during the binding of semantically related information.

On the other hand, our findings are contrary to the linguistic information maintenance theory (Lewis & Bastiaansen, 2015). They proposed that any changes in the processing of the linguistic input (i.e. violations of semantics or syntax) lead to greater beta power desynchronization. However, it is important to note here that in our experimental design we do not induce semantic binding violations but rather compare semantic binding vs. no semantic binding. This may partially explain why the current effects differ from some previous empirical studies. For example, Luo et al., (2010) showed that semantically incongruous sentences elicited a reduced beta power 0 to 200 ms post presentation of the critical word (compared to congruous sentences). The time window of the beta modulation of our results (i.e., immediately after the target word onset) is similar to Luo et al. (2010) indicating that this effect may be related to the anticipatory and prediction processes involving binding of the second word. Furthermore, the direction of the beta power modulation is opposing to our findings as during this time window the young adults displayed a reduced beta power increase in the semantic binding condition (equivalent to the semantically congruous sentences) compared to the no binding condition (similar but not equivalent to the semantically incongruous sentences). Additionally, Lewis et al. (2017), also found a greater beta desynchronization in the incoherent condition (compared to coherent condition) when investigating the effect of semantic coherence at a local sentence level, when using short stories. However, it is important to highlight here that this semantic coherence manipulation was only effective at significantly modulating the beta power at the last sentence presentation (in a story with 4 sentences). Lastly, Wang, Jensen, et al., (2012) found that sentences ending in anomalous words induced a beta power decrease (in the same time window as the N400) over the left temporal areas. These beta modulations significantly correlated with the N400: a larger beta power decrease was associated with smaller N400 amplitudes. The authors suggested that the role of beta oscillations within language comprehension is complex, but it is evident that beta oscillations are involved in semantic unification of items into the wider phrase or sentence context. At this point, the reasons for our somewhat conflicting findings remain unclear. Although we do not know exactly how the current results relate to previous findings, it is crucial to communicate them as this will allow for theories to be updated and developed further.

Compared to the young adults, the binding signature in the healthy older adult group was different, and moreover delayed by about 500 ms. Healthy older adults elicited a smaller beta decrease in the semantic binding condition (vs. no semantic binding). This semantic binding effect in healthy older adults is consistent with Meltzer et al., (2017) who observed a greater magnitude of 8-30Hz event-related desynchronization for word lists (no semantic binding condition) compared to sentences (semantic binding condition) in older adults. However, they also observed this effect in young adults, and therefore it may be surprising that we do not see this condition difference in the young group in an equivalent time window. Furthermore, the binding effect in healthy older adults is in line with the maintenance theory by Lewis and Bastiaansen (2015) and its supporting empirical evidence. The no semantic binding condition led to a greater low beta power decrease ~500 ms after the onset of the target word compared to the semantic binding condition as the language comprehension system detected the need for change. This binding effect in the healthy older adults follows a similar timing pattern as Luo et al. (2010), whereas the young adults did not show any significant differences between the two conditions (binding vs. no binding), implying that surprisingly the language comprehension system did not detect any requirements for change in processing. If the early beta power modulation (during the onset of the second word) in young adults is related to the anticipatory/ prediction activation, the system does not need to ‘listen out’ for any changes later on as it has already predicted the linguistic outcome. It is possible that healthy older adults are sensitive to the requirement for the system to change its processing (more so than young adults) as they were not able to anticipate or predict the incoming binding during the onset of the second word. Though, this is only speculation. This late binding effect in the beta band in healthy older adults is not fully in line with the maintenance hypothesis (Lewis & Bastiaansen, 2015), as we see a beta decrease in both semantic binding and no binding conditions (just a smaller one than in the no semantic binding).

The binding signature we see for healthy older adults is not only delayed but also in the opposite direction to that of the young adults. The finding of an inverse effect between a young and older adult age group is similar to some previous studies. For example, Beese et al., (2019) in an auditory sentence comprehension study tested whether oscillatory power differed across age groups when comparing correctly and incorrectly encoded sentences. They reported that young adults displayed a negative effect (later remembered vs later not remembered sentences produced an alpha decrease). This effect was attenuated in the middle aged adults and completely inversed in older adults (later remembered vs later not remembered sentences produced an alpha increase). The authors attributed this alpha band effect to a shift from cortical disinhibition to inhibition during sentence encoding. Additionally, Poulisse et al., (2020) also found an inverse condition effect between older and young adults in a syntactic binding context (in a two-word phrase paradigm). However, their inverse effect was contrary to our findings (i.e. they found that the syntactic binding effect was associated with a larger alpha power increase in young adults, and a smaller alpha power increase amongst the older adults). Although this pattern is the opposite of our results, our paradigm manipulated semantic and not syntactic binding. Also, it is important to note that Poulisse et al., (2020) observed this inverse condition effect between the groups in a much later time window (0.6-1.05sec) after the phrase presentation compared to our study (0-0.3sec).

As changes in the beta frequencies echo the role of language-related binding (Weiss & Mueller, 2012) we summarize that the semantic binding signature (reflected in the beta frequency) occurred *during* the presentation of the target word (which is when the semantic binding process takes place) in the young adults and was delayed by ~500ms in the healthy older group. However, these conclusions need to be treated with caution, as the interpretations of our results are speculative.

### Suggestions for future research

We found different neural signatures in oscillatory power for young vs. healthy older adults in the present study. Differing functional neural patterns in healthy older adults are commonly interpreted as being compensatory (e.g., Cabeza et al., 2002). However, the term compensation should be reserved for differing neural patterns which are contributing meaningfully to performance (Cabeza et al., 2018; Grady, 2012). The present study was not designed to relate changes in brain function to language performance. Ideally, future studies would incorporate a trial-by-trial semantic comprehension performance measure and characterize which changes in network dynamics are predictive of successful language performance. Investigating the direct relationship between age-related functional neural changes and behavioural performance would be a particularly interesting avenue for future research, and be a necessary step in answering the fundamental question of how the ageing brain adapts to structural decline and reorganises its mechanistic functioning to support language comprehension.

In the present study, we used an experimental paradigm that focused on semantic binding, while minimizing contributions of the ability to make predictions. However, natural language comprehension does rely to some extent on actively making predictions. Making predictions gives language processing a head start (Kuperberg & Jaeger, 2016). Previous studies on young adults have found an alpha power decrease prior to the onset of predictable words (Rommers et al., 2017; Wang et al., 2018), suggesting that in young adults anticipatory binding processes are initiated prior to predictable words being presented. A number of ERP studies have demonstrated that there are age-related changes in the ability to engage prediction mechanisms. Age-related changes in processing predictable sentence endings were evident by the lack of a frontal positivity effect for older (compared to young) adults (Wlotko et al., 2012). Also, N400 amplitudes suggest that older adults do not use the sentence context to pre-activate semantic features of predictable words (Federmeier et al., 2002). Future studies could extend on these findings and examine the age-related changes in oscillatory dynamics supporting the use of predictions during semantic binding.

Lastly, future research should incorporate individual differences measures (and thus, larger sample sizes) to assess which non-linguistic cognitive resources and brain structure properties support the implementation of age-related changes in functional neural characteristics (Peelle, 2019). A quantitative shift in capacity constraints (e.g. higher working memory) can qualitatively impact on the way language is processed, for example, making older adults more able to use contextual semantic information or predictions. Moreover, sufficiently flexible cognitive resources can work together to circumvent structural decline and support functional adaptations, maintaining successful language and communication performance.

## Summary

Healthy older adults have a different oscillatory signature for semantic binding compared to young adults: young adults elicit an early semantic binding signature, around the target word presentation, in the form of a smaller low-beta increase during semantic binding (compared to no semantic binding). On the other hand, healthy older adults display a semantic binding signature ~500 ms later, with a smaller low/high-beta decrease in the semantic binding condition (compared to no semantic binding). Our findings are in line with previous literature suggesting that older adults do extract and make use of contextual semantic information, but there are differences (compared to young adults) with respect to when and how this happens (Federmeier et al., 2003; Federmeier & Kutas, 2005; Stine-Morrow et al., 1999; Wlotko & Federmeier, 2012).

## Data availability

Stimuli and data are available here: https://osf.io/f8grv/

## Authorship Contributions

KS and AM conceptualised the study. RM collected the data and analysed the data. AM and KS supervised data analyses. RM and KS wrote the manuscript. All authors edited the manuscript.

## Acknowledgments

We gratefully acknowledge the help of Denise Clissett, the coordinator of the Patient and Lifespan Cognition Database, for her help with participant recruitment.

