## Supplementary material for "How the healthy ageing brain supports semantic binding during language comprehension"

### Supplementary materials

#### TFR of power between conditions (lexical vs no lexical retrieval & semantic binding vs no semantic binding) for each group (healthy older adults and young adults) separately

Within each age group separately, non-parametric cluster based permutation tests were carried out to compare the time-frequency power changes between the experimental conditions. The experimental manipulations involved: lexical retrieval (real words) vs no lexical retrieval (letter strings); semantic binding (surrounding second word in real word-phrases, including plausible (e.g. swift horse) and implausible (e.g. barking horse) word-phrases) vs no semantic binding (surrounding second word in letter string - real word phrases (e.g. swrfew horse)). The analyses were carried out using the 0 to 3.2sec time window (where 0 is the onset of word 1 and 1.8sec is the onset of word 2), and the pre-defined frequency bands theta (4-7Hz), alpha (8-14Hz), low beta (15-20Hz), and high beta (20-25Hz). We refer to the effects that occurred following word one onset but prior to word two onset as 'lexical retrieval' effects. The effects that arose during or post word two onset are referred to as 'semantic binding' effects. The supplementary Figure 1 displays the condition effects within older healthy adults (A) and young adults (B).

##### *Lexical retrieval within healthy older adults*

There was a significantly smaller theta increase in the lexical retrieval condition compared to the non-lexical retrieval condition ($p = .02$) with a corresponding cluster spanning from 0.3 to 0.65 sec post word one onset, maximal over right occipital and left central electrodes. Secondly, there was a significantly greater theta increase in the lexical retrieval condition compared to the non-lexical condition ($p = .004$). The observed cluster spanned from 0.8 to 1.35 sec and was maximal over bilateral occipital and central channels.

Furthermore, a cluster in the observed data was found in the alpha band. The lexical condition elicited significantly greater alpha suppression compared to the no lexical retrieval condition ($p = .012$). The observed cluster began around the word 1 onset until 0.55 sec and was maximal over the bilateral parietal-central electrodes. An additional cluster in the alpha range extended from 0.6 to 1.75 sec. The cluster-based permutation testing indicated this cluster to show a significantly greater alpha increase over the occipital and central electrodes in the lexical retrieval condition vs no lexical retrieval ($p < .001$).

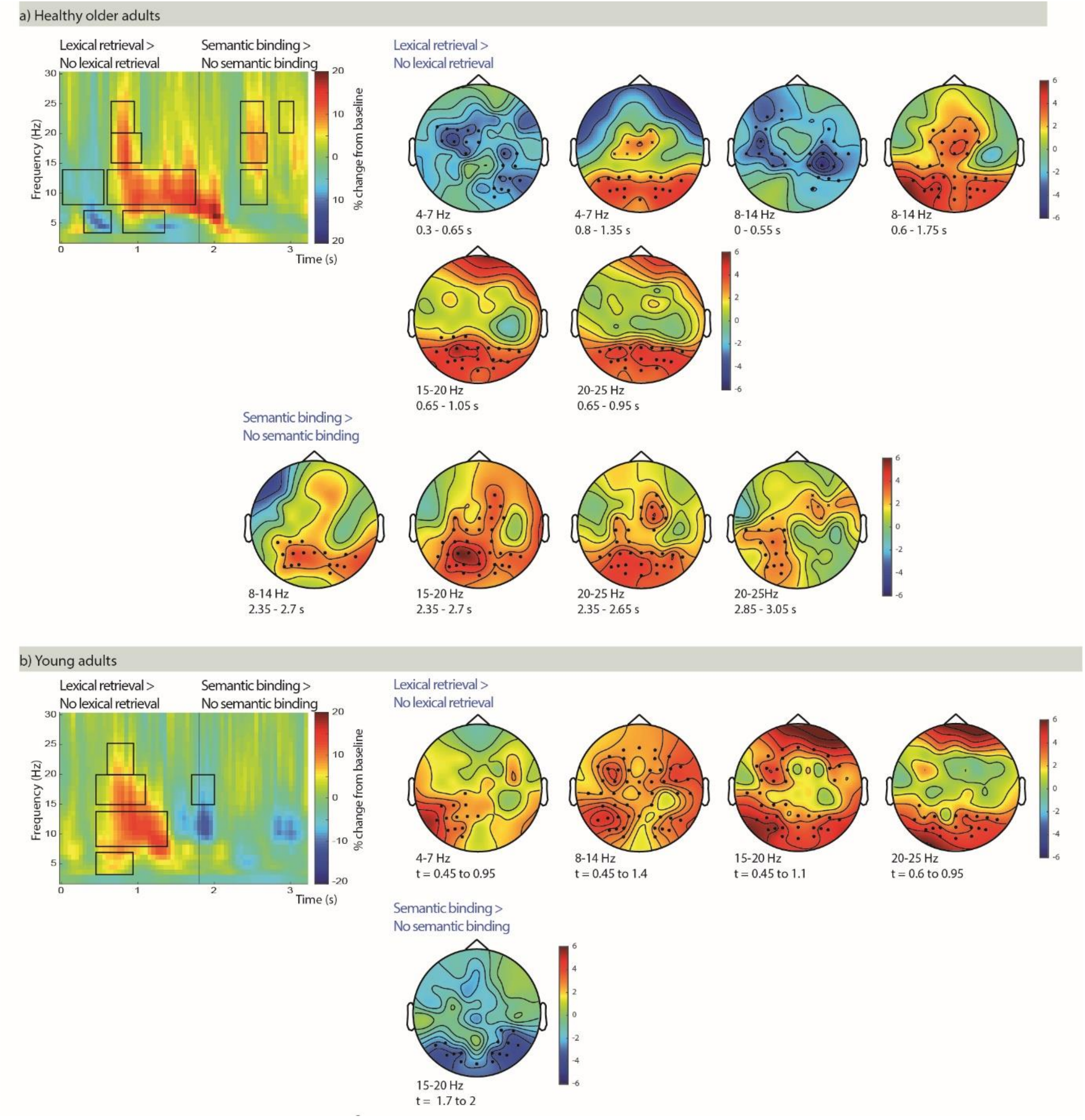


*Supplementary figure 1:* TFRs of power for lexical retrieval/semantic binding minus no lexical retrieval/no binding, in (A) the healthy older adults, and (B) the young adults. Head plots are illustrating the clusters of electrodes that show the most pronounced mean condition difference within each group. Black rectangles indicate significant condition differences ($p<0.05$, cluster corrected).

Furthermore, a cluster was found in the low-beta band. The observed cluster extended from 0.6 to 1.05 sec and was most pronounced over the occipital electrodes. The cluster-based permutation tests indicated a significant condition effect ($p = .002$) (i.e. significantly less low-beta suppression in the lexical retrieval condition post word one onset compared to the no lexical retrieval condition).

Similarly, we observed a cluster in the high-beta band that extended from 0.65 to 0.95 sec over the occipital channels. The cluster-based permutation test indicated significantly less high-beta suppression in the lexical retrieval compared to the no lexical retrieval condition (p = .006).

***Lexical retrieval within young adults***

The cluster-based permutation tests indicated that there were several significant effects of condition. A cluster in the observed data was found in the theta band that extended from 0.45 to 0.95 sec. There was a significantly greater theta power increase (p = .02), maximal over left temporal and parietal channels in the lexical retrieval condition compared to the no lexical retrieval condition.

We also observed a cluster in the alpha band that spanned from 0.45 to 1.45 sec. The cluster corresponded to a significantly smaller alpha power suppression (p = .004) in the lexical retrieval compared to the no lexical retrieval condition, which was evident across the whole scalp.

Lastly, clusters in the low-beta and high-beta bands were observed. They extended from 0.45 to 1.1 sec maximal over frontal, occipital and left temporal electrodes and from 0.6 to 0.95 sec, maximal over occipital electrodes respectively. The clusters corresponded to significant condition effects indicating less low-beta and high-beta suppression in the lexical retrieval condition compared to the no lexical retrieval condition (p = .002 and p = .002 respectively).

***Semantic binding within healthy older adults***

The cluster-based permutation tests found several significant condition effects. A cluster in the observed data was found in the alpha band that extended from 2.35 to 2.7 sec over occipital and parietal electrodes. There was a significantly reduced alpha decrease (p = .038) in the semantic binding condition compared to the no semantic binding condition.

Moreover, clusters in the observed data were found in the low-beta and high-beta ranges spanning from 2.35 sec and ending around 2.65-2.7 sec over occipital, parietal and central electrodes. There was a significantly smaller low-beta (p = .002) and high-beta (p =.002) in the semantic binding condition compared to the no semantic binding condition.

Further condition effects corresponded to a cluster extending from 2.85 to 3.05 sec in the high-beta (maximal over left occipito-central channels) bands. There was a significantly greater high-beta increase in the semantic binding condition compared to no semantic binding (p =.044).

***Semantic binding within young adults***

The cluster-based permutation tests indicated a significant condition effect. The observed cluster was found in the low-beta band extending around the second word onset (1.7 to 2 sec) and was maximal over the occipital channels. The condition effect revealed a significantly smaller low-beta increase (p = .036) in the semantic binding condition compared to the no semantic binding condition.

**Full report of the sub-sampling results**

In order to rule out that the observed condition effects reported in the main paper could be attributed to the difference in the number of trials between conditions, we re-did our analysis matching the number of trials (i.e. random re-sampling) between conditions. We still were able to observe all our previous effects when matching the trial numbers between conditions. We report the results of the re-sampled data below (see also supplementary Figure 2).

**TFR of power between conditions (lexical vs no lexical retrieval & semantic binding vs no semantic binding) for each group (healthy older adults and young adults) separately with the re-sampled data set**

***Lexical retrieval within healthy older adults***

There was a significantly smaller theta increase in the lexical retrieval condition compared to the non-lexical retrieval condition (p < .001) with a corresponding cluster spanning from 0 to 0.6 sec post word one onset, evident across the whole scalp. Secondly, there was a significantly greater theta increase in the lexical retrieval condition compared to the non-lexical condition (p = .009). The observed cluster spanned from 0.8 to 1.35 sec and was maximal over bilateral occipital and central channels.

Furthermore, a cluster in the observed data was found in the alpha band. The lexical condition elicited significantly greater alpha suppression compared to the no lexical retrieval condition (p = .02). The observed cluster began around the word 1 onset until 0.55 sec and was evident across the whole scalp. An additional cluster in the alpha range extended from 0.6 to 1.7 sec. The cluster-based permutation testing indicated this cluster to show a significantly greater alpha increase over the occipital and central electrodes in the lexical retrieval condition vs no lexical retrieval ($p < .001$).

Furthermore, a cluster was found in the low-beta band. The observed cluster extended from 0.65 to 1.05 sec and was most pronounced over the occipital electrodes. The cluster-based permutation tests indicated a significant condition effect ($p < .001$) (i.e. significantly less low-beta suppression in the lexical retrieval condition post word one onset compared to the no lexical retrieval condition).

Similarly, we observed a cluster in the high-beta band that extended from 0.65 to 0.85 sec over the occipital channels. The cluster-based permutation test indicated significantly less high-beta suppression in the lexical retrieval compared to the no lexical retrieval condition ($p = .002$).

***Lexical retrieval within young adults***

The cluster-based permutation tests indicated that there were several significant effects of condition. A cluster in the observed data was found in the theta band that extended from 0.35 to 0.9 sec. There was a significantly greater theta power increase ($p = .007$), maximal over left temporal and parietal channels in the lexical retrieval condition compared to the no lexical retrieval condition.

We also observed a cluster in the alpha band that spanned from 0.45 to 1.4 sec. The cluster corresponded to a significantly smaller alpha power suppression ($p < .001$) in the lexical retrieval compared to the no lexical retrieval condition, which was evident across the whole scalp.

Lastly, clusters in the low-beta and high-beta bands were observed. They extended from 0.5 to 1.15 sec maximal over frontal, occipital and left temporal electrodes and from 0.5 to 1.15 sec, maximal over occipital electrodes respectively. The clusters corresponded to significant condition effects indicating less low-beta and high-beta suppression in the lexical retrieval condition compared to the no lexical retrieval condition ($p < .001$ for both effects).

***Semantic binding within healthy older adults***

The cluster-based permutation tests found several significant condition effects. Firstly, there was a significantly smaller theta increase in the semantic binding condition compared to the no binding condition ($p = .003$) with a corresponding cluster spanning from 1.95 to 2.4 sec, evident across the whole scalp. Furthermore, a cluster in the observed data was found in the alpha band that extended from 2.35 to 2.85 sec over occipital and parietal electrodes. There was a significantly reduced alpha decrease ($p = .011$) in the semantic binding condition compared to the no semantic binding condition.

Moreover, clusters in the observed data were found in the low-beta and high-beta ranges spanning from 2.35 sec and ending around 2.6-2.75 sec over occipital, parietal and central electrodes. There was a significantly smaller low-beta ($p < .001$) and high-beta ($p =.002$) in the semantic binding condition compared to the no semantic binding condition.

Further condition effects corresponded to a cluster extending from 2.85 to 3.05 sec in the high-beta (maximal over left occipito-central channels) bands. There was a significantly greater high-beta increase in the semantic binding condition compared to no semantic binding ($p =.013$).

***Semantic binding within young adults***

The cluster-based permutation tests indicated significant condition effects. The observed clusters were found in the alpha, the low- and high-beta bands extending around the second word onset (1.7 to 2 sec) and were maximal over the occipital channels. The condition effects revealed significantly smaller alpha ($p = .048$), low-beta ($p = .018$), and high-beta ($p = .017$) increases in the semantic binding condition compared to the no semantic binding condition.

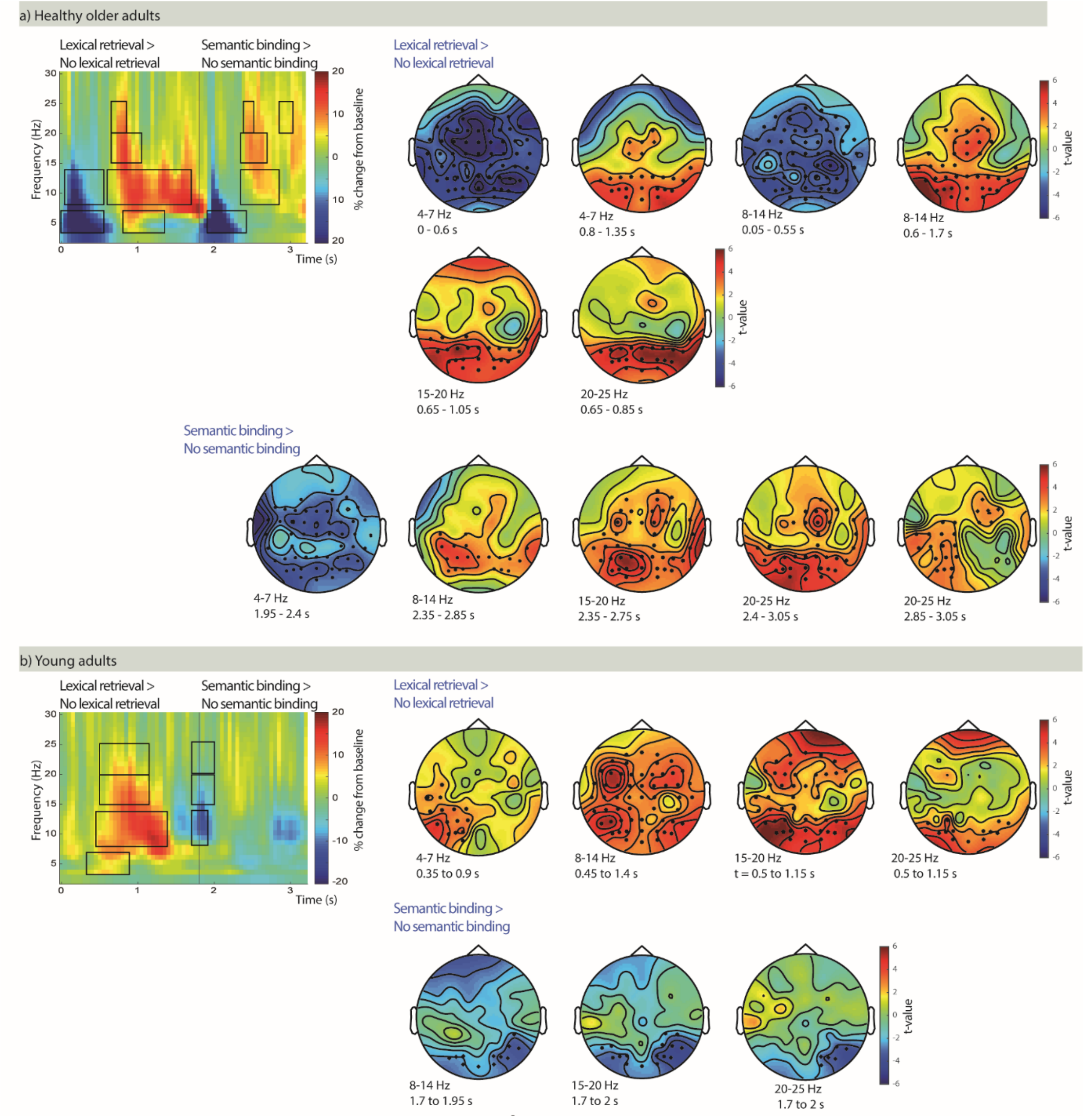


*Supplementary figure 2:* TFRs of power for lexical retrieval/semantic binding minus no lexical retrieval/no binding with re-sampled data, in (A) the healthy older adults, and (B) the young adults. Head plots are illustrating the clusters of electrodes that show the most pronounced mean condition difference within each group. Black rectangles indicate significant condition differences ($p<0.05$, cluster corrected).
